# Breeding the phenylketonuria mouse: novel dietary regimen enables breeding of female C57BL/6J mice homozygous for the *Pah^enu2^* mutation without fostering

**DOI:** 10.1101/2022.12.12.520145

**Authors:** Katherine E. Durrer, Viviana J. Mancilla, Michael S. Allen

## Abstract

The *Pah^enu2^* mutation in C57BL/6J mice is a well characterized model for studying phenylketonuria (PKU), with *Pah^enu2^* homozygotes displaying heightened blood phenylalanine and other characteristics of PKU. *Pah^enu2^* homozygous females do not successfully rear young on any conventional diet due to their disease status. The most commonly used successful breeding strategy is crossing *Pah^enu2^* heterozygous females with homozygous mutant males; producing litters of 50% homozygous and 50% heterozygous animals. In many treatment studies the heterozygous pups produced are not useable, but add to experimental costs and total animals used. To this end our lab created a dietary regimen with reduced phenylalanine and increased large neutral amino acid content, enabling homozygous mating. Phenylalanine levels in homozygous females and males on the new diet are significantly lower than that of homozygous females and males on traditional diets (*p* = 1.35×10^-4^ and *p* = 1.5×10^-5^ respectively). Litters born to *Pah^enu2^* homozygous mothers on this diet demonstrate no significant difference in litter size compared to litters born to heterozygous mothers (*p* > 0.75). No observable defects were noted in litters born from homozygous crosses. This dietary regimen enables litter production of 100% *Pah^enu2^* homozygous animals for investigators who wish to rapidly expand their *Pah^enu2^* mouse colony size or do not require heterozygous littermate controls in their PKU studies.

## Introduction

The *Pah^enu2^* (phenylalanine hydroxylase ENU mutation 2) mouse model of phenylketonuria (PKU) was created on the BTBR background to facilitate research of PKU as an inborn error of metabolism ^1^. This mutation is a T to C point mutation (T835C missense) in exon 7 of the mouse *Pah* gene on chromosome 10 resulting in a phenylalanine to serine substitution at amino acid 263 (F263S) ^2^. Due to the similarities between mice homozygous for the *Pah^enu2^* mutation and human patients with PKU, *Pah^enu2^* homozygous mice have become the gold standard for PKU research. Although several hallmarks of human PKU disease are similar in the *Pah^enu2^* mouse model of PKU, differences between human PKU and mouse PKU are observable; such as the exact amount of phenylalanine in the blood of healthy and PKU individuals (details in discussion).

Shedlovsky et. al. note BTBR homozygous *Pah^enu2^* animals were fertile; however, a very low proportion of pups surviving to weaning age when born to homozygous females even when dams were maintained on a low phenylalanine diet to mediate the hyperphenylalaninemia caused by *Pah^enu2^* homozygosity. An increase in pups surviving to weaning age was only obtained from a homozygous mutant mother when pups from homozygous mutant mothers were fostered by a healthy female, necessitating extra breeding animals, excellent timing of mutant and healthy pregnancies, and euthanasia of pups from the healthy litter ^1^. The inability to breed homozygous females became more pronounced when the mutation was backcrossed to the C57BL/6J background. It is common knowledge in the PKU research community that *Pah^enu2^* homozygous females on the C57BL/6J background cannot be bred successfully and multiple investigators can corroborate this (Jackson Laboratories, Dr. C. Harding, and Dr. K. Skvorak personal communications). Females heterozygous for the *Pah^enu2^* mutation do not have hyperphenylalanemia, breed normally, and do not require litter fostering for their pups to survive. As such, researchers using this animal model have maintained their colonies by breeding heterozygous females with *Pah^enu2^* heterozygous or homozygous male mice. These breeding crosses produce 75% and 50% non-PKU affected pups respectively which are frequently unusable in PKU research projects.

Surplus animals of an unusable genotype are problematic in animal number justifications. All researchers must provide justification for the number of animals proposed for an experiment, including foster animals and animals discarded for an unusable genotype. In addition to the increased number of animals used, the surplus of healthy animals created by breeding heterozygous females adds to the cost of each PKU research project pursued. An ideal solution for the PKU research community would enable direct breeding of homozygous *Pah^enu2^* mouse pairs.

To our knowledge no other laboratory has successfully used any of the alternative low phenylalanine diets to enable breeding of homozygous *Pah^enu2^* mice on the C57BL/6J background. We present results of a *Pah^enu2^* homozygous breeding study using an improved dietary formulation and feeding regimen. The diet includes reduced phe content and enhancement with large neural amino acids, and resulted in successful *Pah^enu2^* homozygous breeding with litter sizes matching that of female mice heterozygous for the *Pah^enu2^* mutation.

## Materials and Methods

### Animal Care and Usage University of North Texas Health Science Center

*Pah^enu2^* mutant mice on the C57BL/6J background (C57BL/6J.BTBR-Pah^enu2^) were acquired from Dr. Harding of Oregon Health and Science University and re-derived at Jackson Laboratory (now available as stock number 029218 from the Jackson Laboratory). Animals were bred and maintained at the University of North Texas Health Science Center in the Department of Laboratory and Animal Medicine’s barrier specific pathogen free facility. All procedures were approved by the University of North Texas Health Science Center Institutional Animal Care and Use Committee under protocol number 2014/15-17-A04.

As previously noted by many investigators and Jackson Laboratories, C57BL/6J mice are very sensitive to alterations in their environment during breeding and pup rearing (cage changes, animal care staff changes, water bottle leaks, slamming doors, etc.) ^3^. Precautions were taken to minimize the occurrence of each stressor during breeding. A cycle of 12 hours of light/12 hours of dark, with the light cycle during the day, and temperature of 22-23°C were maintained at all times.

Water and chow were available to animals *ad libitum*. The standard chow used in non-experimental and non-breeding animals was LabDiet 5LG4 (LabDiet, Purina). Breeder diet for heterozygous animals was LabDiet 5058 (LabDiet, Purina). The experimental PKU breeding diet was TD.160344 (Envigo/Teklad, detailed composition sheet in supplemental materials), a diet made with 40% whole ground oats and a total estimated phenylalanine content of 0.21%. PKU animals were fed the experimental diet for a minimum of two weeks prior to breeding. Animals were not used for breeding unless they were a minimum of eight weeks old and typically breeding was not begun until animals were ten weeks old. Breeding was performed harem style, with one male to every two or three females. Females were separated to their own cage upon noticeable pregnancy and continued to be fed the appropriate diet based on maternal genotype. Cages with pregnant mothers were checked daily for pups. Upon separation, each PKU female began to receive supplemental diet pellets (Love Mash by LabDiet, Purina) twice a week. Quantity of supplemental pellets increased with increased protein production burden of the mother, as previous publications indicate a need for at least 0.22% phenylalanine in the diet for growth of PKU animals (Supplemental diet amounts in Table 1). To ensure pups had access to solid food, some chow pellets were left on the cage bottom once the pups reached two and a half to three weeks of age to encourage solid food intake. Our previous experience with PKU pups indicates that a slightly reduced phenylalanine diet assists in more rapid PKU pup growth. As such, once pups were three weeks old, food in the cage was a 1:1 ratio of regular chow and TD.160344 PKU breeding chow until weaning at 4-5 weeks old, consistent with the age of weaning for the wild-type C56BL6/J mouse strain ^4^. Once weaned, PKU animals were maintained on LabDiet 5058 or a 50/50 mixture of TD.160344 and LabDiet 5058 as needed to prepare animals for other experiments.

**Table 1.** Quantity of Love Mash Supplement provided per breeding mother based on breeding stage and litter size.

| Love Mash | Pregnancy/Pups |
| --- | --- |
| 1 gram | Est. 2.5 weeks pregnant |
| 1 gram | 4 or fewer pups < 1 week old |
| 1.5 grams | 5+ pups < 1 week old |
| 1.5 grams | 4 or fewer pups 1-3 weeks old |
| 2 grams | 5+ pups 1-2 weeks old |
| 2.5-3 grams | 5+ pups 2-3 weeks old |

Heterozygous females were bred to produce pups for other experiments in the lab for the twelve recorded litters in this study. The number of infanticide litters was not recorded for heterozygous mothers, but in previous breedings within our colony the litter infanticide rate is typically one litter for every four to seven first time mothers. Of the nineteen homozygous females bred for this study, two dams killed the litter due to known stressors as mentioned above (water bottle leak on day of birth, cage change on day of birth). Two litters were killed for unknown reasons, this is a common problem in this mouse strain ^3^, and is consistent with infanticides by heterozygous females of this strain. The first ten recorded homozygous female litters for this study were females from mothers that had were *Pah^enu2^* heterozygotes, and these first ten homozygous females were bred to enable statistical comparison with the heterozygous litters. The final five homozygous females (and their male counterparts) were from litters born to homozygous mothers, and these animals were bred to confirm fertility of animals born to homozygous mothers.

Quantifying plasma phenylalanine in animals on the new low phe diet was performed by fluorometric assay as previously described ^5^. Plasma phenylalanine determinations for animals on standard chow were obtained from our previous mouse colony data when animals were maintained at the University of North Texas in Denton. At the time, animals were fed Prolab RMH 1800/5LL2 by Lab Diet, which contains 0.79% phe. Food and water were administered *ad libitum*, a 12 hours of light/12 of dark cycle, and temperature of 22-23°C were maintained at all times with all procedures approved by the University of North Texas Institutional Animal Use and Care Committee, protocol 1202-03. Blood/plasma phenylalanine levels from the Denton colony were determined via blood samples collected and analyzed for genotyping and experimental purposes.

### Mouse blood collection

Dried heparin tubes were prepared as previously described ^5^. A sub-mandibular (cheek) puncture provided blood that was collected into a dry heparin coated tube (no more than 125μl of blood per collection). Whole blood was immediately used for PCR genotyping or frozen at −20°C until needed. If blood was collected for the plasma portion, the microcentrifuge tube with whole blood was centrifuged at 1,000×g for 10 minutes to separate plasma from red and white blood cells. The plasma was removed from the centrifuged blood and placed in a fresh microcentrifuge tube for immediate use or stored at −20°C until testing was performed.

### PCR based *Pah* genotyping of mice

As previously described ^5^, HemoKlen Taq kit (New England Biolabs, USA) was used as needed for genotyping the *Pah* alleles following the manufacturers concentrations and reaction conditions for *Pah* PCR product length and primer sequence. Primer sequences for the amplification of *Pah* were *Pah ^enu2^* forward primer (5’-TGCTGCAACCTGGTAATACTGATCC-3’), and *Pah ^enu2^* reverse (5’-GAACATTGGAGCTTGATGGAATCC-3’). Digestion of the 616 base pair product with restriction enzymes BbsI or BsmAI (Thermo Scientific, USA) were performed as directed by the manufacturer. The *Pah ^enu2^* allele remains uncut when digested with BbsI, while the wild type *Pah* allele is cut into nearly identical bands at 300 base pairs (bp) in length. If using BsmAI, the *Pah ^enu2^* allele is cut into fragment sizes of 308, 274, and 34 bp while the wild type *Pah* allele is cut into DNA of 342 and 274 bp.

### Statistical analysis

Statistical analysis was performed using Microsoft Excel. The two-tailed Student’s t-test was used to compare litter sizes of litters born to heterozygous females and litters born to homozygous female. The two-tailed Student’s t-test was also used to compare plasma phe levels on the two different diets. A *p*-value of 0.05 was used for significance testing. All ± numbers indicate standard deviations.

## Results

In a previous facility (University of North Texas, Denton campus), our *Pah^enu2^* homozygous animals were fed a standard rodent chow containing 0.79% phe. On this standard rodent chow animals displayed plasma phe values of 2109 ± 181μM for males (n = 4) and 2603 ± 405μM for females (n = 6). When fed TD.160344 for two weeks or more, plasma phenylalanine for *Pah^enu2^* homozygous males was significantly reduced to 1230 ± 50μM, n = 5, and plasma phenylalanine for homozygous females was significantly reduced to 1311 ± 185μM, n = 3 (*p* = 1.5×10^-5^ for males and *p* = 1.35×10^-4^ for females). Plasma concentrations of other amino acids were not determined. On standard diets, animals homozygous for the *Pah^enu2^* mutation displayed typical hypopigmentation. When fed TD.160344, fur color of homozygous mutant animals became indistinguishable from healthy heterozygous animals.

All litters in this study are the first litter for each female due to increased litter size in later litters for C57BL/6J females. In our facility, females heterozygous for the *Pah^enu2^* mutation fed the standard mouse chow and bred to males heterozygous or homozygous for the *Pah^enu2^* mutation produced a mean litter size of 5.1± 1.8 pups at weaning age (n = 12). Females and males homozygous for the *Pah^enu2^* mutation fed the diet regimen described in materials and methods produced a mean litter size of 5.2 ± 1.8 pups at weaning age (n =15). No significant difference in litter size was found using a two-tailed student’s t-test (*p* = 0.80). Similarly, weaning age of homozygous mutant pups was uniform, with no alteration in weaning age based on maternal genotype.

Due to the previously mentioned constraint of infanticide of handled pups (a C57BL/6J trait), direct measurements of young pups were not possible. However, pups observed through the clear cages had no observable difference in body or head size in the days immediately after birth. Similarly no animals born to PKU affected females demonstrated microcephaly or hydrocephaly at any age (both of which are readily visible defects in older pups). Behavior of PKU pups born to PKU mothers was not noticeably different from PKU pups born to carrier mothers. Dissections were not performed to determine a presence or absence of cardiovascular defects.

Consistent with the C57BL/6J mouse strain, some litters born to first time homozygous mothers were lost shortly after birth and are not included in the statistics above ^3^. Two litters born to homozygous mothers were lost without a known stressor, and two litters were lost when a known stressor occurred (one litter lost due to a water bottle leak on the day of birth, and one litter lost due to a cage change on the day of birth). A loss of two litters without a known stressor over nineteen total litters is slightly less than or equal to the typical litter loss rate observed in previous colony maintenance of this mouse strain and indicates typical breeding performance of the C57BL/6J mouse.

To ensure animals from homozygous crosses were healthy enough to breed, males and females from homozygous crosses were bred to each other (five of the fifteen total homozygous litters described herein). The mean litter size at weaning for these crosses was 5.0 pups per litter, consistent with the mean litter size of 5.2 pups of all fifteen litters born to homozygous dams.

## Discussion

In order to fully understand the data, it is important to note the differences in blood phenylalanine levels in humans and mice. Healthy and PKU mice exhibit higher levels of phenylalanine than their human counterparts. Healthy human blood phenylalanine is typically 35-76 μM, while classical PKU patients exhibit phenylalanine levels of 1200μM and higher ^6^. Healthy mice (wild type C57BL/6J or carrier animals) in our colony exhibit blood phenylalanine levels between 155 and 250 μM (n =20) on a 0.79% phenylalanine diet. As with human patients, the phenylalanine level in the blood of a PKU mouse is greatly influenced by total dietary phenylalanine intake. When our PKU animals were maintained on a low/moderate protein diet (Prolab RMH 1800/5LL2 by LabDiet) blood phenylalanine levels were 2109 ± 181μM for males (n = 4) and 2603 ± 405μM for females (n = 6). This gender discrepancy of higher blood phenylalanine in female animals with PKU is normal for this model (Dr. K. Skvorak, and Dr. C Harding, personal communications).

In mice it is not possible to bring PKU blood phenylalanine levels down to match the levels of healthy controls when ingesting the minimum phenylalanine required for growth ^7^. In fact, PKU mice on the minimum phenylalanine diet had blood phenylalanine levels approximately 14 times higher than healthy controls ^8^. In mice the required dietary phenylalanine intake is estimated at 0.22% when amino acid composition of said synthetic diet was performed ^9^. When previously feeding our adult (not growing) mice a fully synthetic diet of 0.20% phenylalanine, blood phenylalanine of 881 ± 95 μM (n = 3) was observed in male PKU mice ^10^, consistent with an inability to bring blood phe levels to that of control animals while providing the animals with the minimum phenylalanine needed for survival. On our new diet, male animals displayed blood phenylalanine levels of 1230 ± 50μM, (n = 5). This indicated our new low phenylalanine breeding diet was nearly as effective at lowering blood phenylalanine as the previously described minimum phenylalanine diet for PKU animals.

To create the PKU breeding diet, we incorporated observations from our previous dietary studies in these animals. One undesirable characteristic from previous diets was the greatly reduced animal fecal production and very different fecal texture, likely caused by the reduced fiber content of the synthetic diet (stools on this diet were between diarrhea and a regular stool, unpublished data). Similar to the dislike of fully synthetic protein supplements by PKU patients, animals strongly disliked the fully synthetic low phenylalanine chow used in Denton despite added sugar. It became clear for taste and fiber content, a low phenylalanine chow with a large proportion of naturally sourced ingredient/s would be desirable. Simply substituting the protein source for a protein/peptide low in phenylalanine, such as glycomacropeptide, would not provide the fiber desired nor did we feel would it replace enough of the total chow components to significantly improve taste.

We also felt it was important to incorporate aspects of typical breeding diets when creating our PKU breeding diet. Many traditional rodent breeding diets contain a high proportion of oats. Whole oats provided a reasonable fiber source to address fecal/gastrointestinal concerns and whole oats provided a lower phenylalanine to protein ratio than many other natural foods. In our initial feedings with the new diet, we saw an improvement in fecal quantity and quality when compared to our previous fully synthetic diets for other studies (data unpublished). Furthermore the mice did not reject the new diet, and willingly ate TD.160344.

As previously noted ^3^, a common problem in the C57BL/6J mouse strain is infanticide of the entire litter by a notable proportion of first-time mothers. Furthermore, it is well known within the PKU research community that homozygous *Pah^enu2^* females on the C57BL/6J background fed a standard diet will always kill or abandon birthed litters (Jackson Laboratory, Dr. C. Harding, and Dr. K. Skvorak, personal communications). In fact, mouse studies into the effects of PKU on the offspring of homozygous mothers (maternal PKU) required euthanasia of the pregnant PKU female in order to study the pups born to PKU mothers ^11^. Homozygous crosses using our new dietary regimen did not result in a higher rate of litters killed by first-time mothers; in fact the loss of only two of nineteen litters to homozygous mothers from unknown causes is as low or lower than we would expect for a C57BL/6J based mouse colony.

Another concern when breeding previously unbreedable phenotypes is the health of the offspring for breeding and research studies. Maternal PKU occurs when a mother with PKU is untreated or undertreated during pregnancy ^6, 11^. Maternal PKU results in an increase of several birth defects such as intrauterine growth restriction, microcephaly, neurological defects, and congenital heart defects in human patients ^6, 11^. Although dissections were not performed on the animals to examine cardiovascular patterning, none of the maternal PKU effects listed above were seen in offspring born to female PKU mice when fed our new diet. Additionally pups from homozygous crosses were fertile and produced litter sizes comparable with other crosses. Weaning age of homozygous mutant pups was uniform in all litters, and was not altered by maternal phenotype.

In conclusion, we have created a dietary regimen that enables breeding of female mice homozygous for the *Pah^enu2^* mutation. Although this diet may not be sufficient for animals as a standalone feed indefinitely, implementation of this dietary regimen for the breeding of homozygous *Pah^enu2^* mice should increase efficiency and productivity of a *Pah^enu2^* mouse colony, while reducing costs associated with unwanted heterozygous offspring for PKU research.

## Abbreviations and acronyms

PKU: Phenylketonuria
phe: phenylalanine
*Pah*: phenylalanine hydroxylase gene

## Acknowledgements

We would like to thank the National PKU Alliance for their support of this work through grant (MSA) and fellowship (KD) funding. VM was supported in part by the National Institute Of General Medical Sciences of the National Institutes of Health under Award Number R25GM125587. The content of this work is solely the responsibility of the authors and does not necessarily represent the official views of the National Institutes of Health.

